# A SARS-CoV-2 spike binding DNA aptamer that inhibits pseudovirus infection in vitro by an RBD independent mechanism

**DOI:** 10.1101/2020.12.23.424171

**Authors:** Anton Schmitz, Anna Weber, Mehtap Bayin, Stefan Breuers, Volkmar Fieberg, Michael Famulok, Günter Mayer

## Abstract

The receptor binding domain (RBD) of the spike glycoprotein of the coronavirus SARS-CoV-2 (CoV2-S) binds to the human angiotensin converting enzyme 2 (ACE2) representing the initial contact point for leveraging the infection cascade. We used an automated selection process and identified an aptamer that specifically interacts with CoV2-S. The aptamer does not bind to the RBD of CoV2-S and does not block the interaction of CoV2-S with ACE2. Notwithstanding, infection studies revealed potent and specific inhibition of pseudoviral infection by the aptamer. The present study opens up new vistas in developing SARS-CoV2 infection inhibitors, independent of blocking the ACE2 interaction of the virus and harnesses aptamers as potential drug candidates and tools to disentangle hitherto inaccessible infection modalities, which is of particular interest in light of the increasing number of escape mutants that are currently being reported.

## Introduction

The coronavirus SARS-CoV-2 binds via its spike protein (CoV2-S) to the extracellular domain of the human angiotensin-converting enzyme 2 (ACE2) initiating the entry process into target cells. CoV2-S is a trimeric, highly glycosylated class I fusion protein. It binds to ACE2 via the receptor binding domain (RBD) of its S1 subunit.^1^ The trimeric spike exists in a closed form which does not interact with ACE2 and in an open form where one RBD is in the so-called ‘up’ conformation exposing the ACE2 binding site.^2,3^ Upon RBD binding to ACE2 the interaction between the S1 and S2 subunits is weakened allowing S2 to undergo substantial structural rearrangements to finally fuse the virus with the host cell membrane.^2,3^ The important role of the RBD for viral infectivity is underlined by the analyses of neutralizing antibodies from sera of human re-convalescents, which reveal binding of these antibodies to RBD.^4,5^ Consequently, almost all published neutralizing antibodies developed for therapeutic use target RBD, including humanized monoclonal antibodies^6^, antibodies cloned from human B cells^7-9^ and single-chain camelid antibodies.^10,11^ However, mutations in RBD of CoV2-S can cause RBD-targeted antibodies ineffectual while the virus’s interaction with ACE2 remains unchanged or even found improved.^12^ To address this limitation, additional inhibitors of viral infection and a different mode of action, e.g., by targeting other domains of CoV2-S are highly desired but of limited availability.

Against this backdrop, we here report on a DNA aptamer with a different modality of inhibiting viral infection. As the aptamer does not interact with RBD, it does not interfere with the binding of CoV2-S to ACE2. Regardless, the aptamer inhibits viral infection, exemplified by employing a CoV2-S pseudotyped virus and an ACE2 expressing cell line. These findings demonstrate that viral infection can be inhibited independent of targeting RBD and suggest that inhibition could be possible despite the virus has already bound to cells. The results open the path to inhibitors of SARS-CoV-2 infection with hitherto inaccessible modes of action.

## RESULTS

### Selection and characterization of CoV2-S binding aptamers

To identify single-stranded (ss)DNA aptamers that bind to CoV2-S we employed an automated selection procedure^13^. The trimerized His-tagged extracellular domain of CoV2-S, stabilized in the prefusion conformation, was expressed and purified from HEK293 cells^14,15^ and immobilized for the selection on magnetic beads. After twelve selection cycles (**Supporting Fig. 1a**) the enriched ssDNA libraries were analyzed for improved CoV2-S binding by flow cytometry using cy5-labelled ssDNA and CoV2-S immobilized on magnetic particles (**Fig. 1a**). These experiments revealed an increased fluorescence signal of the ssDNA from selection cycle 12 in the presence of CoV2-S (**Fig. 1a**). No interaction was observed when particles without CoV2-S or particles modified with His6-Erk2 or His6-dectin-1 were used, indicating specificity of the enriched ssDNA library. In contrast, the ssDNA library from selection cycle 1 did not show interaction with the particles, independent of their modification state (**Fig. 1a**). The enriched DNA populations were subjected to next-generation sequencing (NGS), in which 10^6^ to10^7^ sequences were analyzed per selection cycle (**Supporting Fig. 1b**). This analysis revealed a strong decrease in the number of unique DNA sequences, starting from selection cycle 4 and levelling between 10% to 5% of unique DNA sequences in selection cycle 7 to 12 (**Fig. 1b**). Likewise, the distribution of nucleotides within the initial random region changed significantly throughout the course of selection, in which guanine is the most frequently enriched nucleotide (**Supporting Fig. 1c**). These data reveal a strong enrichment of DNA sequences, which is further supported by the occurrence of sequences with high copy numbers, e.g., > 100.000 per sequence starting from the DNA populations from selection cycle 5 onwards (**Fig. 1c**). Further in-depth population analysis revealed the occurrence of sequence families, termed family 8, 13, 22, 29, and 30 (**Fig. 1d, Supporting Fig. 1d, Supporting Table 1**). Whereas the frequency of sequences belonging to family 8 started to enrich from cycle 8 onwards, all other families were observed in the DNA populations from selection cycles 4 to 6, having maximum frequency between selection cycles 7 to 10 and declined afterwards (**Fig. 1d**). We chose representative monoclonal sequences within each family that reflect the enrichment patterns (SP1-7, **Fig. 1e**) and tested them regarding interaction with CoV2-S using flow cytometry. These studies revealed interaction of the family 8 sequences SP5, SP6, SP7 with CoV2-S (**Fig. 1f,g**). All other sequences and a scrambled version of SP5 (SP5sc) as putative non-binding negative control sequence did not interact with the target protein (**Fig. 1f**). SP5, SP6, and SP7 bind with high specificity to CoV2-S; no binding to the isolated RBD, ACE2 or to the spike protein of SARS-CoV (CoV-S) was observed (**Fig. 1h**). Kinetic analysis by surface plasmon resonance (SPR) of the interaction of CoV2-S with 5’-biotinylated aptamer variants immobilized on streptavidin coated sensor surfaces show high affinity binding to CoV2-S, with dissociation constants (K_D_) between 9 and 21 nanomolar (**Tab. 1, Supporting Fig. 2a,b**). All aptamers revealed comparable *K*_*D*_ values at 37°C vs. 25°C (**Tab. 1**). A qualitative assay^16^ to determine the impact of the 5’-modifications on the aptamers CoV2-S binding properties revealed only minor influence of the 5’-cy5-, 5’-biotin-, or 5’-hydroxyl labels (**Supporting Fig. 2c**). SP5 showed a slightly decreased binding in the hydroxyl state whereas binding of SP6 to CoV2-S was found to increase by ∼50% in the unmodified state as compared to the 5’-cy5-modified version (**Supporting Fig. 2c**). The interaction properties of SP7 appeared to be independent of 5’-modifications (**Supporting Fig. 2c**).

**Table 1.** Kinetic properties of the aptamers SP5, SP6 and SP7 measured by surface plasmon resonance.

| Sequence | Condition | $k_a (10^4 \text{M}^{-1} \text{s}^{-1})$ | $k_d (10^4 \text{s}^{-1})$ | $K_D (\text{nM})$ |
| --- | --- | --- | --- | --- |
| SP5 | 37°C | $1 \pm 0.2$ | $37 \pm 36$ | $9.2 \pm 7.9$ |
| SP6 | 37°C | $2.1 \pm 0.6$ | $4.5 \pm 1.9$ | $21 \pm 4.6$ |
| SP7 | 37°C | $1.5 \pm 0.4$ | $2.9 \pm 1.2$ | $18.9 \pm 5.5$ |
| SP5 | 25°C | $1.19 \pm 0.1$ | $1.74 \pm 0.01$ | $14.7 \pm 0.8$ |
| SP6 | 25°C | $2.5 \pm 0.3$ | $3.4 \pm 0.2$ | $13.9 \pm 0.6$ |
| SP7 | 25°C | $1.3 \pm 0.2$ | $1.7 \pm 0.7$ | $13.1 \pm 3.8$ |

**Figure 1:**
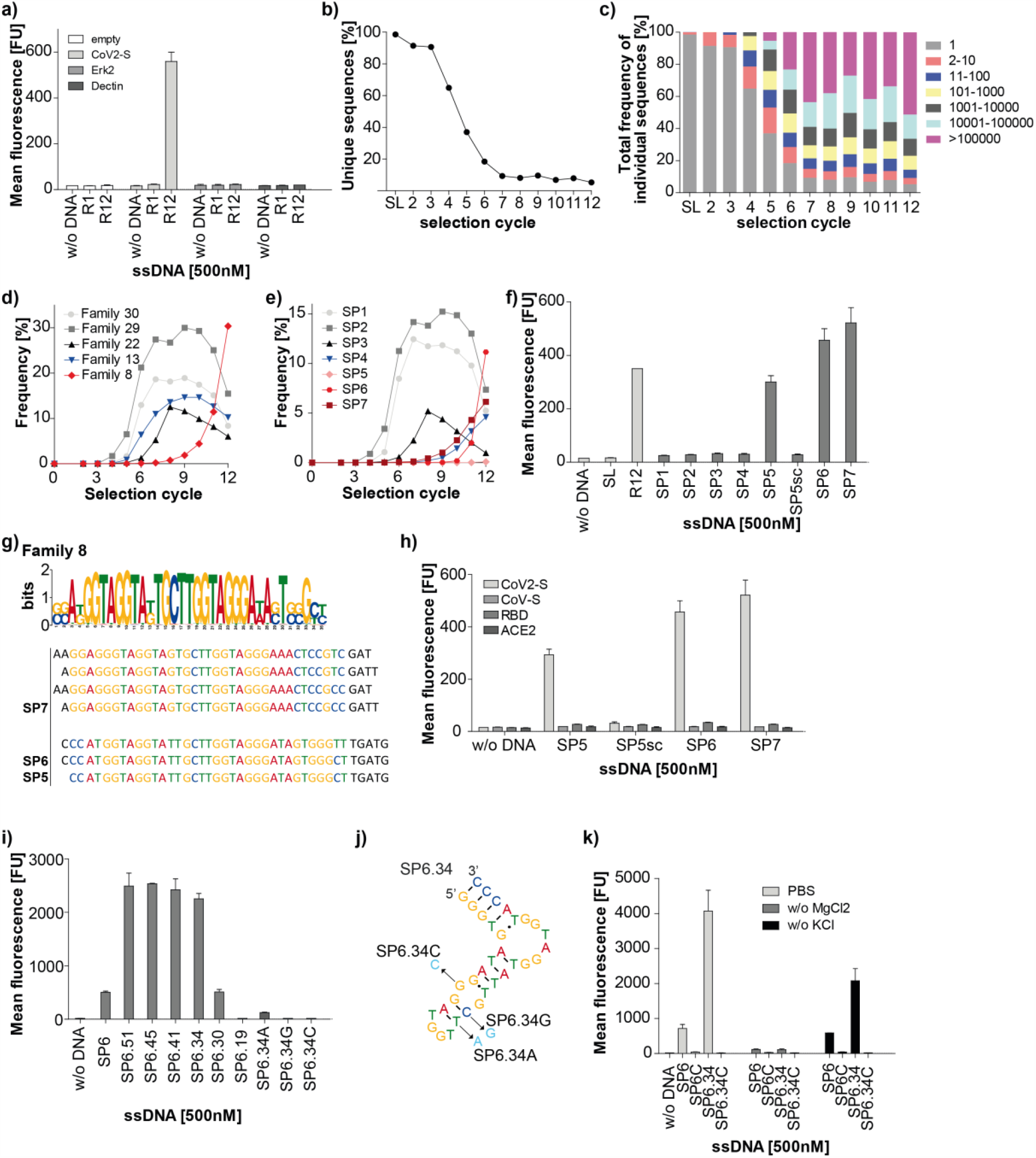
Selection of DNA aptamers binding to CoV2-S. **a)** Interaction analysis of the enriched DNA library from selection cycle 1 (R1) and 12 (R12) in respect to empty beads, CoV2-S, Erk2 and Dectin. **b)** Amount of unique sequences in the DNA populations from selection cycle 1-12 and the starting library (SL). c) Fraction of sequences in the DNA population from selection cycle 1-12 and the starting library (SL) sharing the indicated copy numbers. **d)** Frequency of sequences throughout the DNA population from selection cycles 0-12 belonging to the sequence families 8, 13, 22, 29, or 30. See also Supporting Fig. 1d. **e)** Frequency of representative sequences belonging to one of the families from (**d**). SP1 (Fam 30), SP2 (Fam 29), SP3 (Fam 22), SP4 (Fam 13), SP5-7 (Fam 8). **f)** Interaction analysis of aptamers SP1-7, the starting library (SL) and DNA from selection cycle 12 (R12) with CoV2-S. SP5sc: scrambled control sequence with identical nucleotides as SP5 but with different primary structure. g) Sequence motif of family 8 and assignment of aptamers SP5-7. **h)** Interaction analysis of the scrambled sequence SP5sc and aptamers SP5-7 with CoV2-S, RBD, ACE2, and CoV-S. **i)** Interaction analysis of SP6 and shortened variants and defined single point mutants thereof (**j**). **k)** Interaction analysis of SP6, SP6.34 and the respective control aptamers SP6C (see supporting Fig. 2d) and SP6.34C (**j**) with CoV2-S in the absence and presence of Mg^2+^-ions or K^+^-ions. a,f, h,i, and k: N=2, mean +/- SD.

SP6 as a representative of the family 8 was chosen for further analysis (**Fig. 1g**). Based on secondary structure predictions (**Supporting Fig. 2d**), the aptamer was initially truncated, yielding SP6.51, and analyzed by flow cytometry for CoV2-S binding. Interestingly, SP6.51 showed strongly improved binding compared to the parental SP6 aptamer (**Fig. 1i**). Further truncation of SP6, yielding variants with 45 nucleotides (nt, SP6.45), 41 nt (SP6.41), or 34 nt (SP6.34) maintained the elevated binding properties. When SP6 was truncated to 30 nt (SP6.30) binding fell back to the level of the original SP6 aptamer whereas the interaction with CoV2-S was entirely lost by removing additional 11nt (SP6.19, **Fig. 1i, Supporting Fig. 2d**). Moreover, based on the secondary structure prediction of SP6.34, we investigated the interaction of the point mutants of the minimal aptamer variant SP6.34, namely SP6.34A, SP6.34G and SP6.34C with CoV2-S by flow cytometry. These point mutants were chosen to either stabilize (SP6.34C) or destabilize (SP6.34A, SP6.34G) the putative apical stem structure (**Fig. 1j**). However, all point mutants revealed severely diminished binding to CoV2-S, whereas binding of SP6.34A was still detectable (**Fig. 1i**), albeit to a much lesser extent than SP6. Mutating the positions equivalent to SP6.34C in the parental aptamer SP6, yielding SP6C, also abolishes CoV2-S binding (**Fig. 1k**). To conclude the characterization of SP6, the impact of mono- and divalent ions on CoV2-S binding was assessed by flow cytometry of both, the parental and minimal variant of the aptamer. These studies reveal that the binding of SP6 to CoV2-S is sensitive towards the presence of K^+^ and strongly depends on Mg^2+^-ions (**Fig. 1k**). The binding of the parental aptamer SP6 to CoV2-S was maintained in the absence of K^+^-ions, whereas the interaction of the minimal variant SP6.34 was found to be reduced by about 50% compared to its level obtained in PBS (**Fig. 1k**). These data indicate that K^+^-ions are most likely required for supporting structure formation of the aptamer, which is more pronounced in the truncated variant than in the parental full-length aptamer, but not essential for CoV2-S binding.

### Aptamers selected for the RBD do not interact with CoV2-S

We also performed automated selection procedures to target the isolated RBD of Cov2-S (**Supporting Fig. 3**). Conventional automated selection conditions, as applied targeting CoV2-S (**Fig. 1**), resulted in strong overamplification during the PCR step (**Supporting Fig. 3a**), which could be decreased by reducing the amount of target (10% compared to the conventional approach, **Supporting Fig. 3b**) or by adding heparin as a competitor during the incubation step of the selection procedure (**Supporting Fig. 3c**). Interaction analysis of the obtained DNA libraries from the selection cycles in which no or very low overamplification was observed, i.e. cycle 6 of the conventional procedure (**Supporting Fig. 3a**), cycle 9 when less target was used (**Supporting Fig. 3b**), and cycle 8 when heparin was added (**Supporting Fig. 3c**), revealed enrichment of RBD binding species in all selections (**Supporting Fig. 3d**). However, none of the enriched RBD-binding libraries interacted with full-length CoV2-S comprising the complete extracellular domain (**Supporting Fig. 3d**). The starting library, used as negative control, neither bound to RBD nor to CoV2-S, whereas the library enriched for CoV2-S (R12 CoV2-S, **Fig. 1a**), used as positive control, showed binding to CoV2-S as expected (**Supporting Fig. 3d**). Of note, library R12 CoV2-S also revealed interaction with RBD, although to a lesser extent than to CoV2-S (**Supporting Fig. 3d**). Therefore, we decided to use the library R12 CoV2-S to conduct three additional selection cycles (cycles 13-15) enriching for those species that bind predominantly to RBD instead of other domains of CoV2-S, that are presumably targeted by SP5, SP6, and SP7 (**Fig. 1h**). We again used the conventional selection approach (cycles 13-15, **Supporting Fig. 3e**) and a selection variant with less (10%) RBD than in the preceding selection (cycles 13*-15*, **Supporting Fig. 3e**). In both cases overamplification was observed from cycle 13/13* on, although less pronounced as during the *de novo* selection targeting RBD under previously applied selection conditions (**Supporting Fig. 3a**). Both enriched libraries (R15/R15*) showed binding to RBD but no interaction with CoV2-S (**Supporting Fig. 3f**). Next-generation sequencing of the obtained libraries revealed two strongly enriched distinct families (**Supporting Fig. 3g-i, Supporting Tables 2,3**). We chose four representative sequences, RBD1-4, and performed interaction analysis. These experiments were found to be in-line with the observations obtained with the enriched libraries, i.e. the sequences bound to RBD (**Supporting Fig. 3j**) but not CoV2-S (**Supporting Fig. 3k**). Despite RBD4, which was found at elevated copy numbers in selection cycle 6 of the selection targeting CoV2-S but declining thereafter, all RBD binding sequences only increased in copy numbers when the target changed from CoV2-S to RBD in selection cycles 13-15 (**Supporting Fig. 3l**) and 13* to 15* (**Supporting Fig. 3m**). These data indicate that targeting RBD of CoV2-S with DNA libraries, in our hands, was not productive in yielding aptamers interacting with the full-length extracellular domain of CoV2-S protein *in vitro*.

### SP6 inhibits viral infection independent of the interaction of CoV2-S with ACE2

We next performed pulldown experiments to further characterize and verify the interaction of SP6 with CoV2-S (**Fig 2a**). In these experiments, biotinylated SP6 or SP6C were incubated with the respective protein and the complexes were collected by adding streptavidin coated magnetic beads. After washing, the remaining proteins were analyzed by SDS-PAGE and Coomassie staining of the gel. Aliquots were taken and analyzed prior to the incubation with the magnetic beads (input, **Fig. 2a**), from the supernatant after incubation with the magnetic beads (unbound, **Fig. 2a**) and from the bead/aptamer bound fraction (eluate, **Fig. 2a**). SP6 revealed binding to CoV2-S (**Fig. 2a**, eluate, lane 2) but not to CoV-S (eluate, lane 6) nor to ACE2 (eluate, lane 4) or the unrelated control protein Nek7 (eluate, lane 5). In this experiment, SP6C showed weak binding to CoV2-S (**Fig. 2a**, eluate, lane 1). In agreement with the results obtained by flow cytometry (**Fig. 1h**) binding of SP6 to CoV2-S was not reduced even in the presence of a fivefold molar excess of RBD (**Fig.2a**, eluate, lane 3).

**Figure 2:**
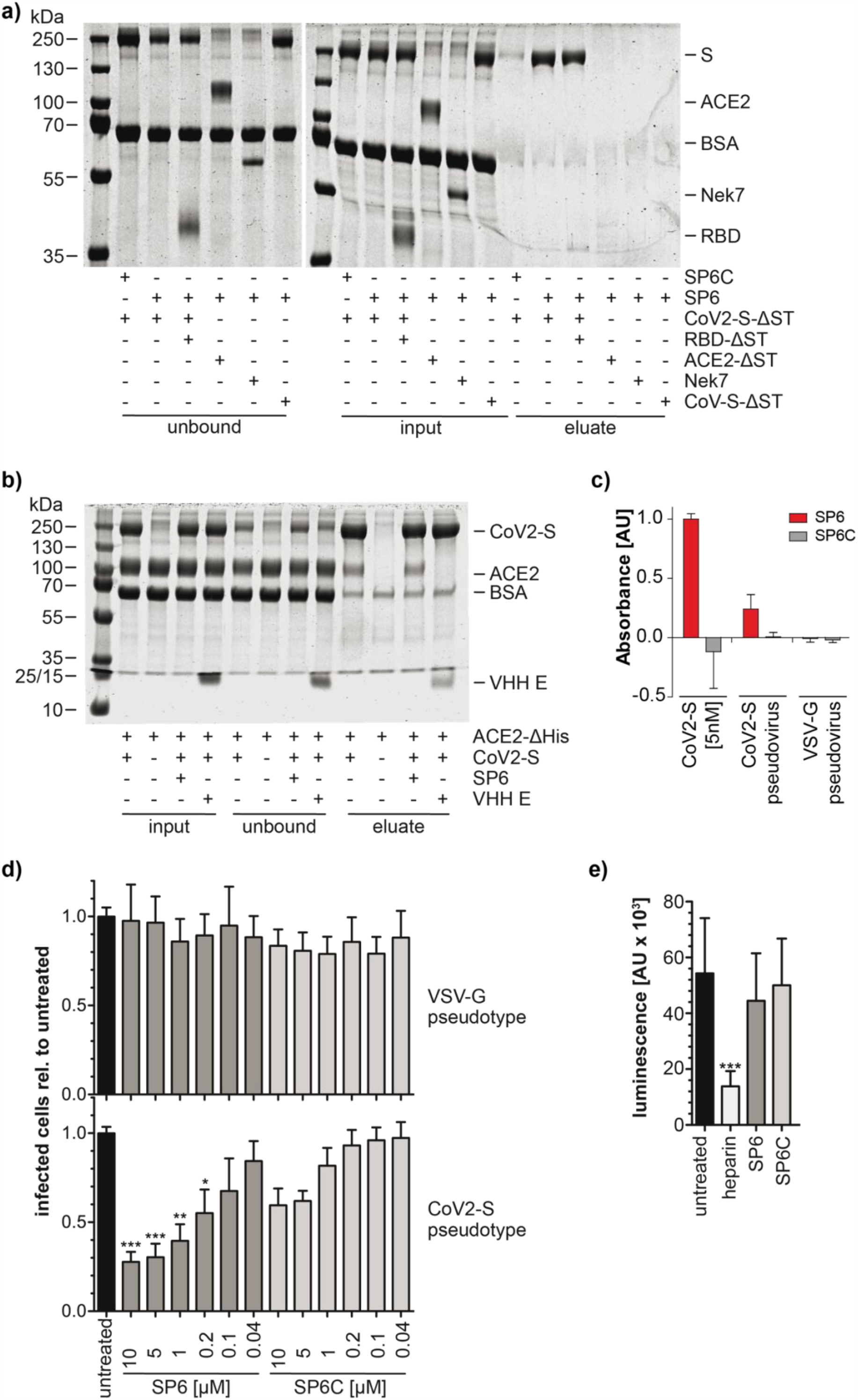
RBD independent inhibition of CoV2-S pseudovirus infection. **a)** Pulldown analysis of SP6 binding specificity. ΔST indicates constructs lacking the StrepTag. **b)** Pulldown analysis of CoV2-S ACE2 interaction. ΔHis indicates lack of His tag. **c)** ELONA of S protein and SARS-CoV-2-S pseudotype virus. **d)** SARS-CoV-2-S pseudovirus infection. n=5, mean +/- SD, *** p<0.001, ** p<0.01, * p<0.05. **e)** SARS-CoV-2-S pseudovirus binding. n=8, mean +/- SD *** p<0.001

As SP6 appears not to interact with the RBD of CoV2-S, we investigated whether SP6 has an impact on the interaction of CoV2-S and ACE2. To this end, His-tagged CoV2-S was pulled by Ni-NTA magnetic beads and the co-pulldown of untagged ACE2 (ACE2ΔHis) was analyzed in the presence or absence of SP6 (**Fig. 2b**). As before, input, unbound and eluate fractions were analyzed by SDS-PAGE and Coomassie staining. Whereas the interaction between CoV2-S and ACE2 was abolished by the RBD-binding control nanobody VHH E (**Fig. 2b**, eluate, lane 4), SP6 did not affect this interaction (eluate, lane 3). Densitometric analysis of the respective bands resulted in ACE2:Cov2-S ratios of 0.18 and 0.16 in the absence (eluate, lane 1) or presence (eluate, lane 3) of SP6, respectively.

Having shown SP6 interacts with CoV2-S without interfering with complex formation with its cellular receptor ACE2, we next studied the impact of SP6 on viral infection. To address this question, we used the established VSV-ΔG*-based pseudotype system^17,18^ and generated Cov2-S and VSV-G pseudotyped virus particles. The interaction of SP6 with the CoV2-S pseudotyped virus was verified by an enzyme-linked oligonucleotide assay (ELONA).^19^ In this experiment, the CoV2-S protein or the CoV2-S pseudotyped virus were captured by a nanobody binding to the RBD of CoV2-S and after washing the bound protein or pseudovirus particles were detected by adding biotinylated SP6, streptavidin-horse radish peroxidase (HRP) conjugates and its substrate 2,2′-Azino-di(3-ethylbenzthiazoline-6-sulfonic acid) (ABTS) (**Supporting Fig. 4**). We observed a concentration dependent increase in signal when SP6 and SP6.34 were used for detection, but not when employing SP6C and SP6.34C (**Supporting Fig. 4a**). Likewise, SP6 but not SPC6C detected the CoV2-S pseudotyped virus. The VSV-G pseudotype was not detected demonstrating the specificity of the assay (**Fig 2c**). Next, ACE2-expressing Vero E6 cells were infected with Cov2-S or VSV-G pseudotyped virus particles, which had been pre-incubated with SP6 or SP6C (**Fig. 2c**). Pseudotype particle numbers were adjusted to result in infection rates between 8 % and 10 % for the aptamer-untreated pseudotypes (**Supporting Fig. 4b,c**). This infection rate was chosen to prevent multiple infections of a single cell precluding reliable measurements. SP6 showed a concentration-dependent reduction of infection of Vero E6 cells by the CoV2-S pseudotype virus (**Fig. 2d, Supporting Fig. 4b,c**). In contrast, the infection of the VSV-G pseudotype was not affected (**Fig. 2d**). These results demonstrate the dependence of the inhibitory effect of SP6 on the presence of CoV2-S on the viral particles and exclude unspecific effects on the infection process of the VSV-G vector. The presence of SP6C also led to some reduction of infection which, however, did not reach statistical significance. The seeming discrepancy to the lack of binding of SP6C to CoV2-S in the binding assay (**Fig. 1k**) or the ELONA (**Fig. 2c**) is explained by the higher concentrations of SP6C used in the infection assay. In addition, unmodified SP6 (as used in the infection assay) shows stronger binding to CoV2-S than the 5’-modified versions (**Supporting Fig. 2c**) and this can also be assumed for SP6C. The slight inhibitory effect of SP6C is in-line with its observed weak interaction with CoV2-S in the pulldown assay (**Fig. 2a**).

Whereas ACE2 is the most important receptor for CoV2-S, at least two co-receptors are known to contribute to CoV2-S binding to target cells, heparan sulfate and neuropilin-1.^20,21^ Therefore, we investigated whether SP6 affected binding of CoV2-S pseudotyped particles to cells even if it did not inhibit CoV2-S binding to ACE2. For this purpose, VSV-ΔG* was pseudotyped with CoV2-S carrying a HiBiT tag at the C-terminus. Vero E6 cells were incubated with these particles and bound virus was quantified by NanoBiT reconstitution (**Fig 2e**). Whereas the known inhibitor heparin^22^ reduced binding of CoV2-S pseudotyped particles, neither SP6 nor SP6C had an effect on binding. These data show that SP6 reduces pseudovirus infection by interfering with a process occurring after binding of the pseudovirus to cells.

## Discussion

In conclusion, we describe the DNA aptamer SP6 binding to CoV2-S and with the potential to inhibit SARS-CoV-2 infection. A remarkable and unexpected feature of SP6 is that its inhibitory effect does not result from interfering with the interaction of CoV2-S with ACE2. This feature distinguishes the mode of action of SP6 from that of CoV2-S targeting antibodies. Currently, the overwhelming majority of these antibodies bind to the RBD of CoV2-S^6-11^ and prevent ACE2 interaction by either directly competing with ACE2 for binding^6-10^ or by stabilizing an ACE2 binding-incompetent conformation^11^. Antibodies not binding to RBD but to the N-terminal domain of CoV2-S have also been shown to prevent interaction of CoV2-S with ACE2 although by an yet unknown mechanism.^10^ To our knowledge, neutralizing antibodies targeting the S2 domain have not yet been described. At present, the molecular mechanism by which SP6 inhibits viral infection is unknown. As the binding of CoV2-S pseudotypes to cells is not affected, we conclude that a step occurring after binding must be impeded. This could involve preventing S2’ cleavage or destabilizing the prefusion conformation of the spike protein. The latter mechanism has been shown to lead to viruses bearing spike proteins in the postfusion conformation and has been reported for an antibody neutralizing SARS-CoV.^23^ This antibody, however, binds to the RBD. We anticipate that the elucidation of the mechanism by which SP6 inhibits infection will provide insight into how CoV2-S triggers fusion of the viral and host cell membranes.

There is an increasing number of currently reported mutations in SARS-CoV-2^9^, among which the most recent example is the apparently faster spreading lineage VUI-202012/01, also named B.1.1.7.^24^ This variant shows several mutations in the RBD resulting in escape of binding to some antibodies. Since more escape mutations in the RBD can be expected to further arise in the future, RBD-independent modalities to prevent infection, as revealed by SP6, are of relevance and need to be investigated.

SP6 might be further optimized to increase potency. For example, homo- or heterovalent multimers could be engineered by combining SP6 with itself or aptamers (or other ligands) binding to different CoV2-S domains, a strategy employed previously to gain very potent thrombin inhibitors.^25,26^ Indeed, di- or trimerization of CoV2-S antibodies has been shown to increase their potency.^11,27^ The automated selection process enables the rapid generation of aptamers, for example for mutated proteins of SARS-CoV-2 lineages that escape treatment regimens by aptamers, antibodies, or other active pharmaceutical ingredients. Likewise, re-selection strategies to adapt SP6 towards mutations are also possible. In addition to these features, aptamers provide means to develop antigen tests, exemplified by the presented SP6-based ELONA data. The ease by which aptamers can be synthesized, their low batch-to-batch variations and long shelf lives predestines SP6 for various diagnostic and treatment options, e.g., as inhalation spray.

## Acknowledgement

This work has been made possible by funds from the Federal Ministry of Education and (BMBF, 01KI20154 to MF and GM) and by the German Research Council (DFG, MA3442/7-1 to GM). We thank Dr. F.-I. Schmidt and Dr. P.-A. König (Core Facility Nanobodies, Medical Faculty, University of Bonn) for providing the RBD nanobody and Prof. J. Schultze for performing NGS (University of Bonn/DZNE). We are grateful to Dr. G. Zimmer (Institute of Virology and Immunology, Mittelhäusern, Switzerland) for providing the VSV-ΔG* and to Prof. J.S. McLellan (The University of Texas at Austin, USA) for providing the plasmid for CoV2-S expression.

## Author contribution

AS developed the cellular binding and partly executed the cellular binding and infection assays, constructed the pseudotype viruses and wrote the manuscript. AW performed sequence analysis of the enriched libraries and conducted interaction analysis of the DNA libraries, aptamers and their variants. MB conducted infection and binding experiments. SB performed the automated selection experiments. VF expressed and purified proteins and performed the pulldown experiments. MF supervised experiments, discussed results and wrote the manuscript. GM conceived and designed the study, supervised and discussed results and wrote the manuscript. All authors have read and commented on the manuscript.

## Supporting Information

### Material and Methods

#### Coupling of SARS-CoV-proteins to Dynabeads His-Tag isolation & pulldown

For immobilization of SARS-CoV-2 proteins, Dynabeads His-Tag Isolation & Pulldown (ThermoFisher) were used. For this purpose, 9.6 nmol of SARS-CoV-proteins, prepared in 1 mL wash/binding buffer (50 mM Sodium-Phosphate, pH 8.0, 300 mM NaCl, 0.01% Tween®-20) were coupled to 100 µL of bead solution (40 mg beads/mL), according to the manufacturer’s protocol. The provided buffer, by the manufacturer, of the bead solution was discarded by separation before coupling, using a DynaMag™-2 Magnet (ThermoFisher). For the coupling reaction, the solution was incubated for 30 min on 4°C, using a Tube Revolver Model D-6050 (neoLab) rotating at a speed of 20 rpm. According to the manufacturer’s protocol, three washing steps with 1 mL wash/binding buffer were carried out, followed by one additional washing step with storing buffer (ssDNA selection = 1.25× PBS; 171.25 mM NaCl (Fisher Scientific), 3.38 mM KCl (Roth), 12.5 mM Na_2_HPO_4_ (Roth), 2.2 mM KH_2_PO_4_ (Roth), pH 7.4; 1 mg/mL Albumin (BSA) Fraction V (pH 7.0) (AppliChem); 3.25 mM MgCl_2_ // 2’fRNA selection = 1.25× PBS; 171.25 mM NaCl (Fisher Scientific), 3.38 mM KCl (Roth), 12.5 mM Na_2_HPO_4_ (Roth), 2.2 mM KH_2_PO_4_ (Roth), pH 7.4; 1 mg/mL Albumin (BSA) Fraction V (pH 7.0) (AppliChem), before resuspending SARS-CoV-2-protein beads in 1 mL of storing buffer. In the particular case of competitor, 0.125 mg/mL Heparin was added to the storing buffer.

#### Automated selection of ssDNA aptamers

For the selection of ssDNA aptamers, the D2 DNA library (5′ – GGG AGA GGA GGG AGA TAG ATA TCA A – N40 – T TTC GTG GAT GCC ACA GGA C − 3′) was used (Ella Biotech GmbH, Martinsried, Germany). For amplifying the library, the following primers were used: forward primer (Cy5-D2 fw) 5′ Cy5 –GGG AGA GGA GGG AGA TAG ATA TCA A – 3′ and reverse primer (Phos-D2 rv) 5′ P – GTC CTG TGG CAT CCA CGA AA – 3′. PCR master mix for amplification reaction (colorless GoTaq® Flexi Buffer (Promega), 2 mM MgCl_2_ (Roth), 0.2 mM dNTP (Genaxxon)). The PCR reaction was performed by using GoTaq® G2 Flexi DNA Polymerase (Promega), including 1 µM of Cy5-D2 fw and Phos-D2 rv primers, in a total reaction volume of 100 µL and the cycling program 30 s 95 °C, 30 s 62 °C, and 30 s 72 °C in a TRobot thermal cycler (Biometra). In the first four selection cycles 18 PCR cycles were used and in all following selection cycles 16. For all steps performed on the TRobot, an arched auto-sealing lid (Bio-Rad) was used to seal the reaction plate. 1 µL GoTaq® G2 Flexi DNA Polymerase (5u/µL, Promega) was added to start the PCR reaction. The automated pipetting steps were performed by a Biomek NX^P^ (Beckman Coulter). The automated selection was started by 0.5 nmol of D2 ssDNA library pipetted to the SARS-CoV-proteins immobilized on Dynabeads His-Tag Isolation & Pulldown (ThermoFisher) and an incubation for 30 min at 37 °C while shaking at a speed of 700 rpm; pipetting up and down every 5 min during incubation. Selection buffer was PBS/3 mM MgCl_2_/0.8 mg/mL BSA (PBS: 137 mM NaCl (Fisher Scientific), 2.7 mM KCl (Roth), 10 mM Na_2_HPO_4_ (Roth), 1.76 mM KH_2_PO_4_ (Roth), pH 7.4). After incubation, the samples were washed two times with 100µL wash buffer (PBS/3 mM MgCl_2_). Washing steps were increased every selection cycle by two more washes until a total of eight washes per selection cycle was reached. Prior to PCR, the bound ssDNA molecules were recovered by incubation in ddH_2_O (TKA Wasseraufbereitungssysteme GmbH) for 5 min at 80 °C. After PCR, a lambda exonuclease (final 20u, ThermoFisher) digestion was performed for 60 min at 37°C to generate ssDNA for the next selection cycle.

#### Agarose gel analysis

Agarose LE (Genaxxon) was used to prepare 4% agarose gels pre-stained with ethidium bromide (Roth). 1 µL 6x DNA Loading Dye (Thermo Scientific) was mixed with 5 µL of dsDNA products and loaded on the gel. As reference, 4 µL of GeneRuler Ultra Low Range DNA Ladder (Thermo Scientific) were loaded. A Genoplex system (VWR) was used for visualization of the stained dsDNA.

#### DNA interaction analysis

For interaction studies of the selected libraries and individual sequences CoV2-S, RBD, CoV-S or ACE2 were immobilized on magnetic particles (ThermoFisher) via a His-tag according to manufacturer’s instructions and stored in 1.25× PBS with 1 mg/ml BSA at 4°C. 4 µl of the bead suspension were incubated with 500 nM DNA in a final volume of 10 µl in PBS with 3 mM MgCl_2_ and 0.8 mg/ml BSA at 37°C in a thermomixer (Eppendorf) and 650 rpm. After incubation, the beads were washed three times with 100 µl PBS/3 mM MgCl_2_. Bound Cy5-labeled sequences were analyzed by flow cytometry counting approximately 20.000 events. Unmodified sequences were analyzed via OliGreen (ThermoFisher) fluorescence. Therefore, bound DNA sequences were recovered in 100 µl ddH_2_O for 5 min at 95°C, cooled down to 4°C and incubated in 200 µl TE buffer (10 mM Tris/HCl pH 7.5, 1 mM ethylenediaminetetraacetic acid) with a 1:500 dilution of OliGreen. Fluorescence intensity was measured on a Tecan Ultra plate reader (Tecan) at excitation and emission wavelengths of λ = 500 nm and λ = 525 nm, respectively.

#### NGS

The starting library and all enriched libraries were analyzed by NGS using the Illumina HiSeq1500 platform. Samples were prepared according to.^28^ Briefly, 12 different index primers were attached to the different libraries via PCR, allowing the analysis of 12 samples in parallel on one lane. PCR products were purified using the NucleoSpin Clean-Up kit (Macherey Nagel), and equal amounts of PCR product of each library were mixed to a final amount of 2 μg DNA. For the hybridization to the flow cell a subsequent adapter ligation step was performed according to the manufacturer’s instructions (TruSeq DNA PCR-Free Sample Preparation Kit LT, Illumina). Samples were purified by agarose gel purification. The NGS data was analyzed using the inhouse AptaNext software and MEME suite.^16^ Five families from cycle 12 were identified and the most abundant sequences were tested in a FACS binding assay. Secondary structures of the aptamers were predicted with Mfold.^29^

#### SPR

Binding affinities of the full-length sequences were assessed by surface plasmon resonance on a BIAcore 3000 instrument (GE Healthcare Europe). All buffers were filtered and degassed prior to use. 50 nM of biotinylated aptamers SP5, SP6 and SP7 (flow cells 2-4) and the control SP5sc (flow cell 1) were immobilized on XanTec SAD chips (XanTec Bioanalytics) with a flow rate of 10 μl/min at 25 °C in 0.5 M NaCl until values of ∼200 response units were reached. The CoV2-S protein (PBS/3 mM MgCl_2_ and 1 mg/ml BSA) was injected at different concentrations of 1, 3.2,10, 32, 100, 200, 316, 700 and 1000 nM for 180 s at 25 °C and 37°C. The dissociation time was 400 s, followed by a regeneration step (0.5% sodium dodecyl sulfate). Data was evaluated using the BIAevaluation 4.1 (Biacore) software: 1:1 binding with drifting baseline.

#### ELONA

33 ng/ml of the RBD-nanobody (kindly provided by P.-A. Albert and F.-I. Schmidt, Core Facility Nanobodies, Medical Faculty, University of Bonn) was immobilized in 20 µl bicarbonate/carbonate buffer (pH 9.6) on 96 halve-well microtiter plates (MICROLON® 600, VWR) at 4°C overnight followed by two washing steps with 100 µl PBS with 0.05% Tween 20. Wells were blocked with PBS including 5% BSA for 1 hour at RT, followed by two washing steps with 100 µl PBS/3 mM MgCl_2_. Afterwards the CoV2-S protein or the pseudovirus (∼15,000 particles) were added in PBS with 3 mM MgCl_2_ and 0.8 mg/ml BSA and incubated in a final volume of 20 µl at RT, followed by two washing steps with PBS/3 mM MgCl_2_. Next, biotinylated DNA aptamers [500 nM] and controls in PBS/3 mM MgCl_2_ and 0.8 mg/ml BSA were added in a final volume of 20 µl at RT, followed by two washing steps with PBS/3 mM MgCl_2_. Streptavidin-HRP (GE Healthcare) in a 1:1000 dilution in PBS/3 mM MgCl_2_ was added (20 µl) at RT, followed by two washing steps with PBS/3 mM MgCl_2_. Finally, 100 µl ABTS (ThermoFisher) were added per well, incubated at RT for 40 min and the absorbance at λ = 405 nm was measured on a Tecan Nanoquant (Tecan).

#### Constructs and plasmids

Plasmids for SARS-CoV-2-Se, SARS-CoV-2-S-HexaPro and SARS-CoV-Se (kindly provided by Jason McLellan, The University of Texas at Austin, USA) code for the prefusion-stabilized ectodomains of the S proteins and carry on the C-terminus a trimerization motif, a HRV 3C cleavage site, 8xHis and TwinStrep tags. ^14^ SARS-CoV-2-S(D614G)-Δ19 codes for the S protein (GenBank NC_045512) containing the D614G mutation and lacking the C-terminal 19 amino acids thus deleting the ER-retention motif and enhancing transport to the plasma membrane. SARS-CoV-2-S(D614G)-Δ19-HiBiT contains the HiBiT tag (Promega) at the very C-terminus. SARSCoV-2-S-RBD codes for amino acids 319 - 591 of the S protein and contains the signal peptide of the S protein on the N-terminus to allow secretion and an HRV 3C cleavage site, 8xHis and TwinStrep tags on the C-terminus. Proteins denoted by ΔHis or ΔST do not contain the respective tag but are otherwise identical to their tag-containing counterparts. ACE2e contains, after cleavage of the signal peptide, amino acids 19 - 615 of human ACE2 (UniProt Q9BYF1), an N-terminal myc tag and a C-terminal HRV 3C cleavage site followed by an 8xHis tag. The myc and ACE2 coding sequence was amplified form pCEP4-myc-ACE2 (addgene #141185).^30^ With the exception of SARS-CoV-2-S(D614G)-Δ19-HiBiT which was cloned into pmCherry-N1 (Takara) replacing mCherry all these proteins were cloned into pCAG which is based on pCAGGS only lacking the SV40 ori of the latter. pET-Sumo-Nek7 contains the full-length human Nek7 (UniProt Q8TDX7) cloned into pET-Sumo (Thermo). All constructs were assembled from PCR-amplified fragments using Q5 DNA Polymerase (NEB) or synthetic genes (Eurofins) except for the pCAG backbone which was linearized by restriction digestion. For assembly the NEBuilder HiFi DNA Assembly Master Mix was used (NEB). Coding sequences of all constructs were verified by Sanger sequencing (Eurofins).

#### Protein expression and purification

With the exception of Nek7 which was expressed in *E. coli* BL21(DE3) proteins were expressed in FreeStyle 293F cells (Thermo). 293F cells at 1×10^6^ cells / ml in FreeStyle 293 Expression Medium (Thermo) were transfected with 1 mg plasmid and 2 mg PEI max (Polysciences) per liter of cells. 3 - 5 days after transfection proteins were purified from the culture medium after removing cells and debris by centrifugation (10 min, 800 g, rt, followed by 30 min, 10000 g, 4 °C). The cleared medium was adjusted to 50 mM HEPES/KOH, pH 7.8 / 300 mM NaCl / 25 mM imidazole and loaded overnight onto a column containing 2 ml Protino Ni-NTA Agarose (Macherey-Nagel) per liter of medium. After washing with 50 mM HEPES/KOH, pH 7.8 / 300 mM NaCl / 25 mM imidazole proteins were eluted in the same buffer containing 1 M imidazole. Eluted proteins were concentrated using Vivaspin Turbo concentrators (Sartorius) and loaded on a Superose 6 column (Cytiva) equilibrated in 20 mM HEPES/KOH, pH 7.8 / 150 mM NaCl to remove aggregated material. Peak fractions were pooled, concentrated, flash-frozen in liquid nitrogen and stored at −80 °C. Purification of proteins lacking the 8xHis tag was done by replacing the Ni-NTA column by a StrepTactin column (IBA). Elution was achieved by 30 mM desthiobiotin. To remove the 8xHis TwinStrep tags from SARSCoV-2-S-RBD and SARS-CoV-Se the proteins were incubated overnight with HRV 3C protease (Thermo) and passed over a Ni-NTA agarose column. The flow-through was concentrated and further purified by size exclusion chromatography as above.

#### Pulldown assays

For aptamer pulldowns 1 µM 5’-biotinylated SP6 or SP6C, respectively, were incubated with 0.5 µM of the indicated proteins (without TwinStrep tag) for 30 min at rt in buffer PB (PBS / 4 mM MgCl_2_ / 2.5 µM BSA). For the RBD competition, 0.5 µM S protein together with 2.5 µM RBD (without 8xHis and TwinStrep tags) were used. An aliquot was removed (input) and 100 µl pre-equilibrated Hydrophilic Streptavidin Magnetic Beads (NEB) were added. After 30 min incubation at rt on an overhead rotator beads were collected on a magnet and an aliquot was removed from the supernatant (unbound). After washing two times with 500 µl PBS / 3 mM MgCl_2_ proteins were eluted by boiling in Lämmli sample buffer (eluate). For spike-ACE2 complex pulldowns 1 µM SARS-CoV2-S-HexaPro and ACE2 (without 8xHis tag) were incubated in the presence of 3 µM SP6 or VHH E, as indicated, in buffer PB. An aliquot was removed (input) and 20 µl pre-equilibrated HisPur Ni-NTA Magnetic Beads (Thermo) were added. After 30 min incubation at rt on an overhead rotator beads were collected on a magnet and an aliquot was removed from the supernatant (unbound). After washing two times with 250 µl buffer PW proteins were eluted with 25 mM HEPES/KOH, pH 7.8 / 150 mM NaCl / 1 M imidazole (eluate). Samples were separated by SDS-PAGE. Coomassie-stained gels were scanned on an Odyssey Sa (Licor).

#### Pseudovirus generation

VSV pseudotypes were generated as published.^31^ Briefly, Hek293T cells transfected with pCAG-SARS-CoV-2-S(D614G)-Δ19 or pcDNA3.1-VSV-G, respectively, were inoculated with VSV-ΔG* (kindly provided by Gert Zimmer, Institute of Virology and Immunology, Mittelhäusern, Switzerland). In VSV-ΔG* the VSV-G open reading frame is replaced by an expression cassette for GFP and firefly luciferase^17^ allowing infected cells to be detected by GFP fluorescence or luciferase activity. After 1 h incubation at 37 °C the inoculum was removed, the cells were washed with DMEM and cultivated for 16 - 18 h in DMEM / 2 % FBS / 30 mM Hepes at 34 °C. The culture medium containing the pseudotyped particles was clarified from cellular debris by centrifugation (800 g, 5 min, rt). Aliquots were flash-frozen in liquid nitrogen and stored at −80 °C.

#### Infection

Vero E6 cells were cultivated in DMEM (Thermo) / 10 % FBS (PAN) at 37 °C and 8 % CO_2_. The day before infection 5×10^4^ cells were plated per well of a 24well plate. Virus particles pseudotyped with SARS-CoV-2-S(D614G)-Δ19 or VSV-G were pre-incubated for 20 min at rt with the indicated agent in DMEM / 3 mM MgCl_2_. The culture medium was removed from the cells and replaced by 150 µl pre-incubated virus (MOI ≈ 0.1). After incubation for 1 h at 37 °C 0.5 ml DMEM / 10 % FBS / 3 mM MgCl_2_ was added and the cells were cultivated for 16 - 18 h. Cells were detached with trypsin, fixed for 20 min at rt with 4 % formaldehyde, pelleted (800 g, 5 min, rt) and resuspended in PBS. Infection rate was determined as percentage of GFP-positive cells by flow cytometry (BectonDickinson). Data analysis was done with FlowJow 10.7.1 (BectonDickinson). Doublets were excluded (for gating see Supporting Fig. 4c). For statistical analysis the non-parametric Kruskal-Wallis test and Dunn’s multiple comparison post-test was used because due to the small sample size of n=5 Gaussian distribution for the values could not be tested. Analysis was performed with Prism 5.0f (GraphPad).

#### Binding

1×10^4^ Vero E6 cells were plated per well of two 96well plates the day before. Virus particles pseudotyped with SARS-CoV-2-S(D614G)-Δ19-HiBiT were pre-incubated for 20 min at rt with the indicated agent in DMEM / 3 mM MgCl_2_. The culture medium was removed from the cells, replaced by 100 µl pre-incubated virus, and the cells were incubated for 1 h at 37 °C. The inoculum was completely removed and 50 µl of Nano-Glo HiBiT Lytic Reagent (Promega), beforehand diluted with an equal volume of PBS, was added. After 15 min incubation at rt luminescence was measured with an Infinite M1000 Pro (Tecan). Statistical analysis was done as above, n=8, each with 3 technical replicates.

#### Overview on sequences of aptamers used in this study (point mutations are depicted in red bold letters)

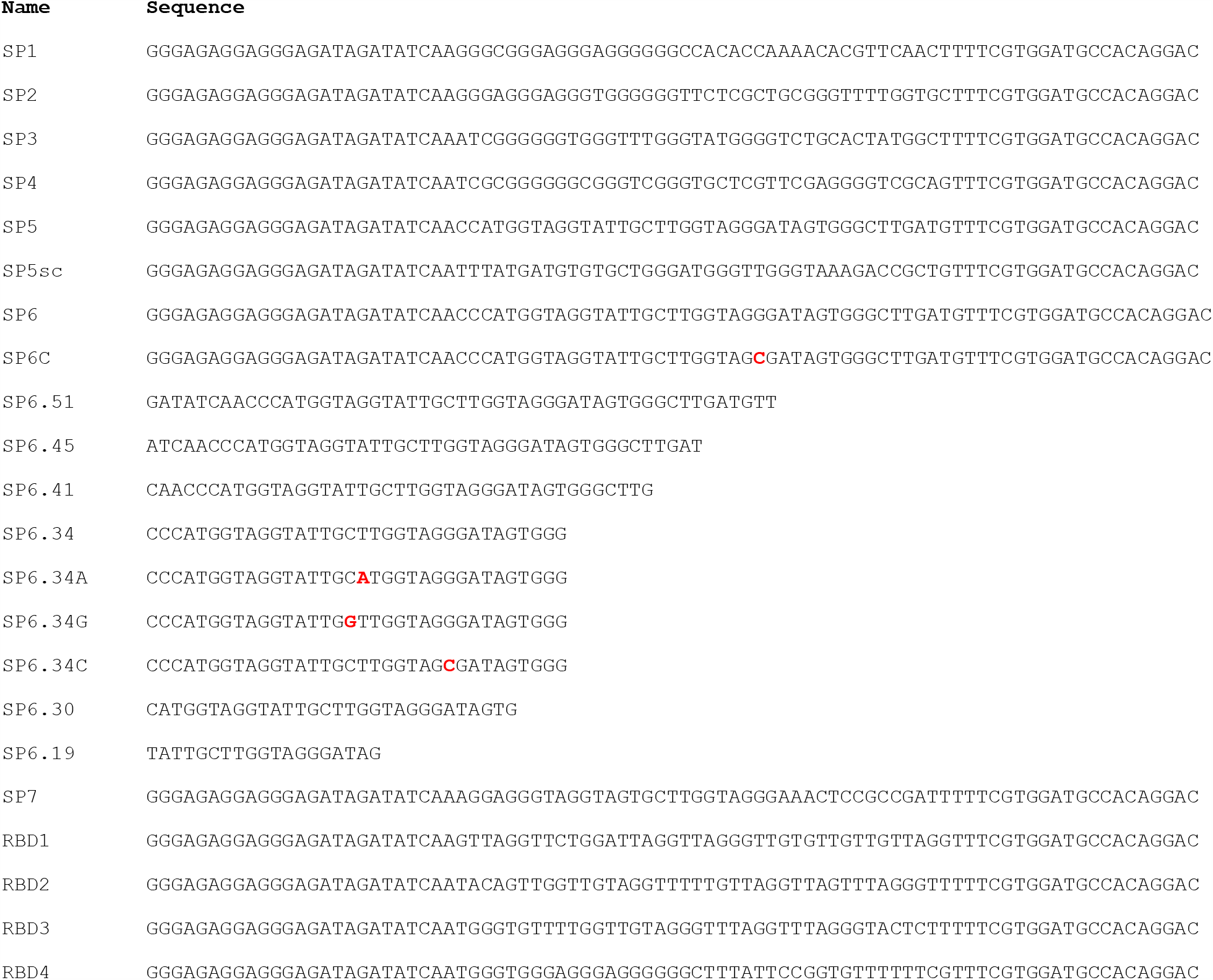

**Supporting Figure 1:**
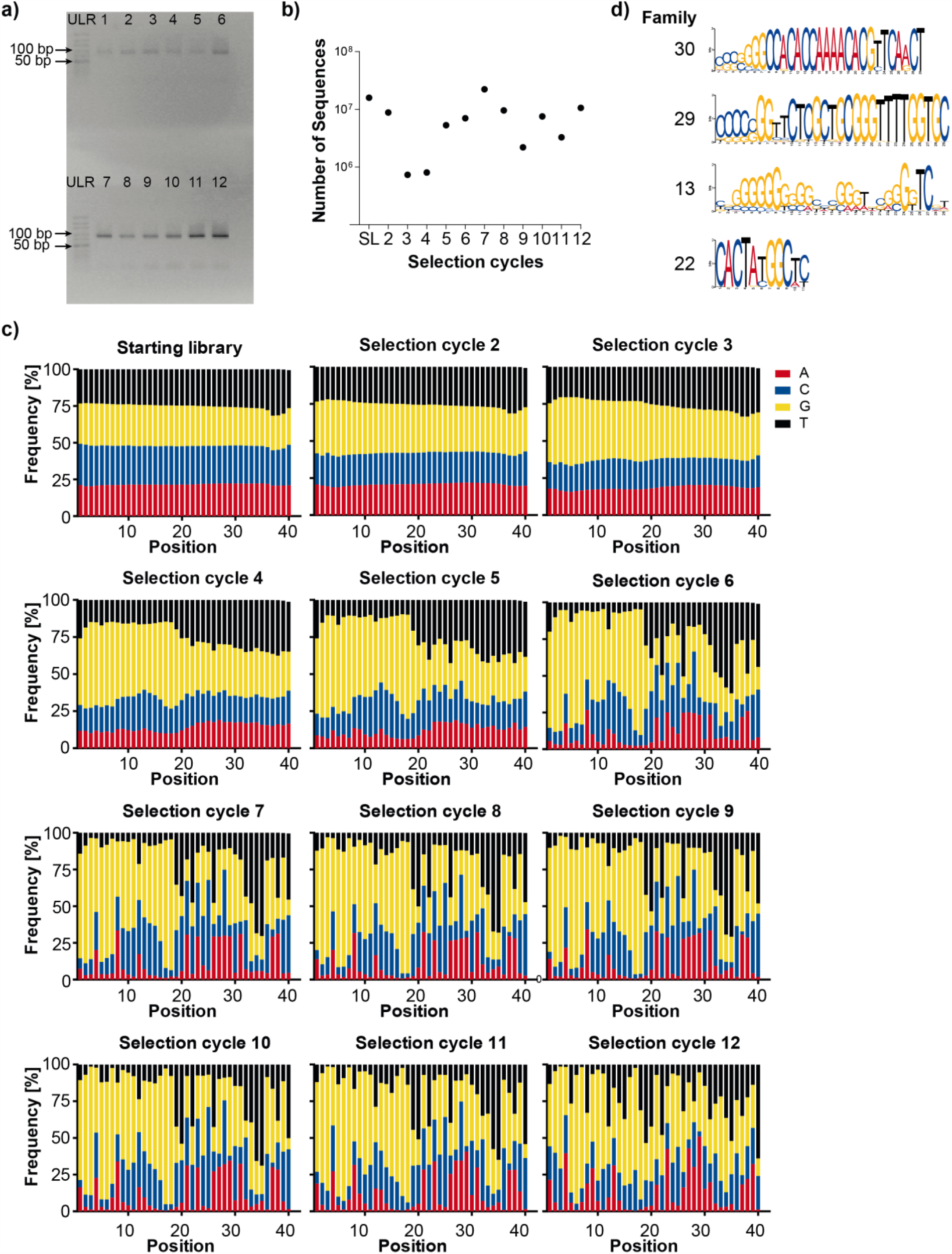
**a)** Agarose gel analysis of the PCR products obtained from the selection cycles 1-12 of the automated selection targeting CoV2-S. ULR: GeneRuler Ultra Low Range DNA Ladder; 1-12 indicate the selection cycle. **b)** Number of analyzed sequences per selection cycle by NGS. **c)** Nucleotide distribution of the random region at the different positions in the starting library and selection cycles 2 to 12. **d)** The predicted families 30, 29, 13, and 22 obtained by NGS analysis based on motif search by MEME.

**Supporting Figure 2:**
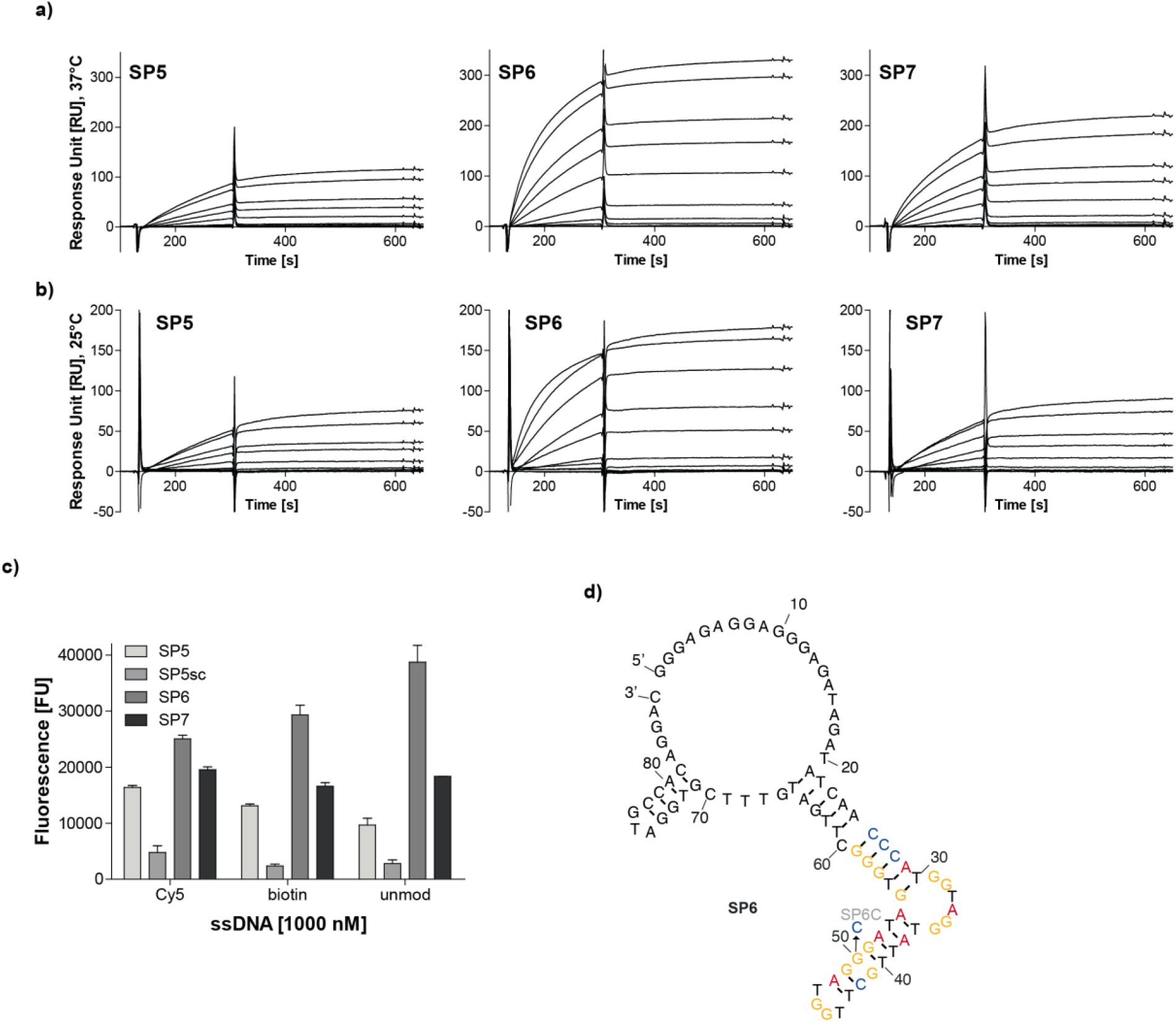
**a,b)** Sensograms of the SPR analysis of biotinylated aptamers SP5, SP6 and SP7 immobilized on a streptavidin chip and the CoV2-S at concentrations between 1-1000 nM at 37°C (**a**) (N = 4) and 25°C (**b**) (N = 2). **c)** OligoGreen-based assay to evaluate the interaction of the aptamers SP5, SP6 and SP7 having different modifications (Cy5, biotin or unmodified) at the 5’-end with CoV2-S. N = 2, mean +/- SD **d)** secondary structure prediction by Mfold of the aptamer SP6. Nucleotides 26 to 59 of the truncated aptamer SP6.34 are highlighted. SP6C depicts the single point mutant G50C.

**Supporting Figure 3:**
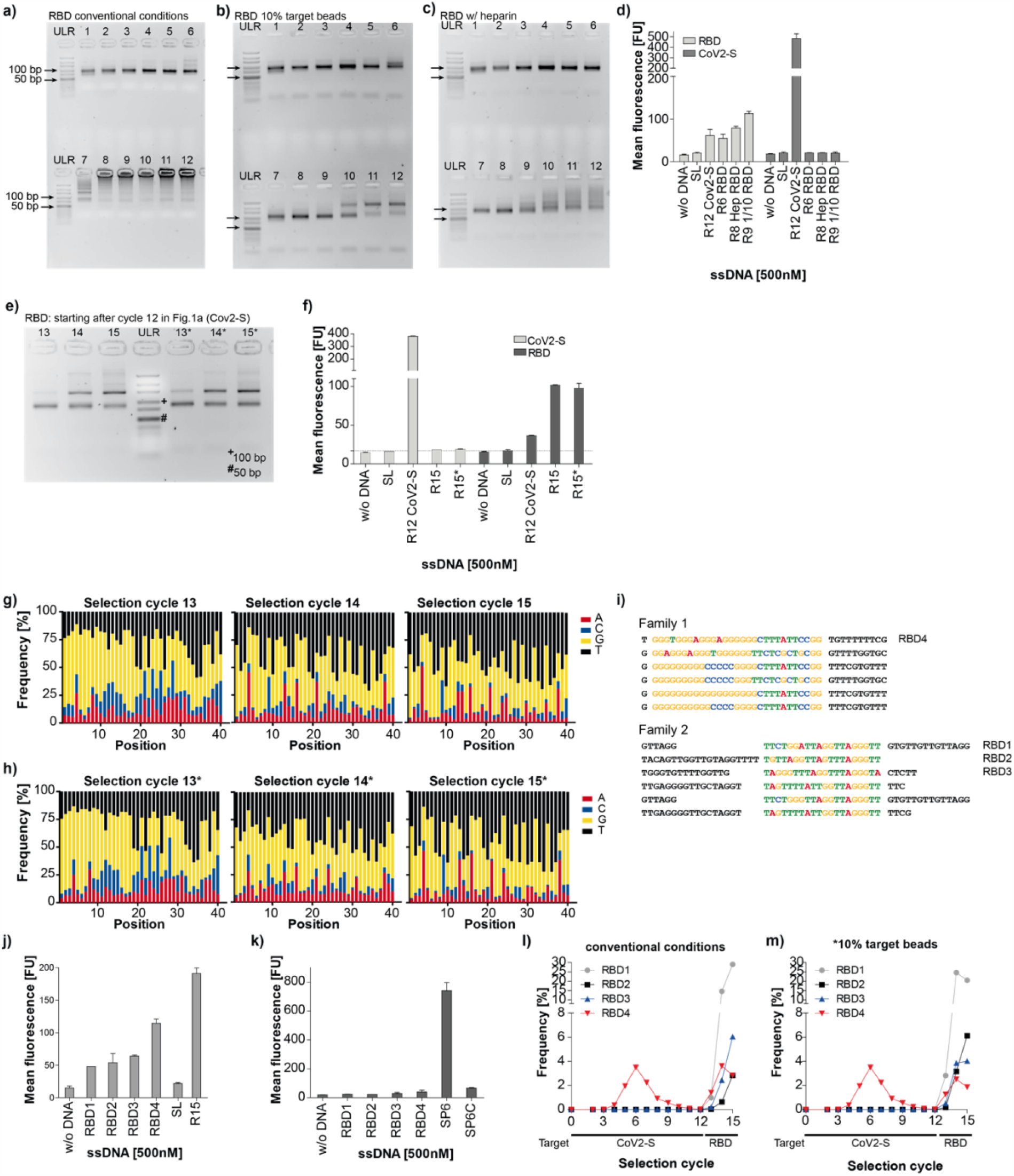
**a-c)** Agarose gel analysis of the PCR products obtained from the selection cycles 1-12 of the automated selections targeting RBD and using conventional selection conditions (**a**), 10% of target (**b**) or heparin as competitor (**c**) during the selection. ULR: GeneRuler Ultra Low Range DNA Ladder; 1-12 indicate the selection cycle. **d**) Interaction analysis of the enriched libraries (**a**)-(**c**) binding to RBD and CoV2-S. SL: Starting library, R12CoV2-S (library enriched for CoV2-S as in **Fig. 1a**). **e**) Agarose gel analysis of the PCR products obtained from the re-selection cycles 13-115 and 13*-15* of the automated selections targeting RBD and starting from the library of selection cycle 12 enriched for CoV2-S (Fig. 1a). ULR: GeneRuler Ultra Low Range DNA Ladder; 13-15 indicate the selection cycles using conventional selection conditions; 13*-15* indicate selection cycles using 10% of the target during the selection. **f**) Interaction analysis of the enriched libraries from (**e**) binding to RBD and CoV2-S. SL: Starting library, R12CoV2-S (library enriched for CoV2-S as in **Fig. 1a**). **g,h**) Nucleotide distribution at the different positions of the random region of the libraries obtained from the re-selection experiments targeting RBD shown in (**e,f**). **i**) The predicted families 1 and 2 obtained by NGS analysis based on motif search by MEME from libraries 15* and 15. Interaction of aptamers RBD1-4 with RBD (**j**) and CoV2-S (**k**). SL: Starting library; R15: as R15 in **f**. Enrichment profiles of RBD1-4 during the selection targeting CoV2-S (selection cycles 1-12, **l,m**) and the two different re-selection conditions, conventional conditions (cycles 13-15, **l**) and with less target protein (cycles 13-15, **m**). d, f, j, k: N = 2, mean +/- SD.

**Supporting Figure 4:**
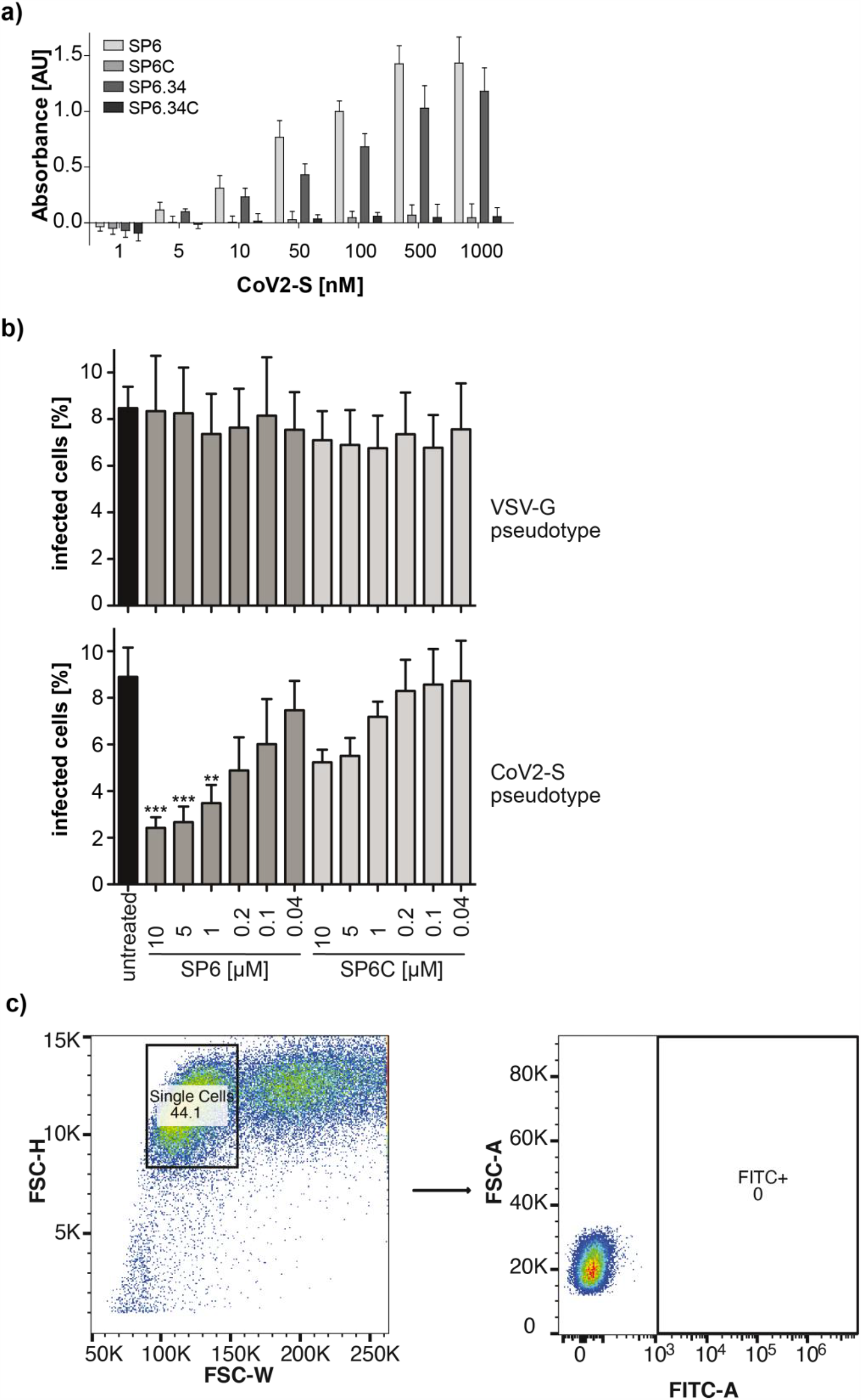
**a)** ELONA assay in sandwich format using a RBD-binding nanobody to capture CoV2-S and the indicated biotinylated oligonucleotide sequence [500 nM] to detect bound CoV2-S. N = 3, error represents standard deviation **b)** SARS-CoV-2-S pseudovirus infection. n=5, *** p<0.001, ** p<0.01. **c)** Gating strategy for quantifying GFP expressing cells by flow cytometry.

